# Inconsistent Subthalamic Beta Expression in the Local Field Potential amid In- and Anti-Phasic Neuronal Bursts

**DOI:** 10.64898/2025.12.16.694607

**Authors:** Maximilian Scherer, Nina Wiedemann, Kai Bötzel, Matthias Löhle, Thomas Kriesen, Daniel Cantré, Jan Hinnerk Mehrkens, René Reese, Thomas Koeglsperger

## Abstract

**Background:** Elevated beta (12-32 Hz) activity in the local field potential (LFP) of the subthalamic nucleus (STN) is a hallmark of Parkinson’s disease and closely tied to synchronized spiking. Albeit the biomarker’s prevalence has not been independently and reliably assessed, it is employed to guide deep brain stimulation (DBS) lead implantation. Likewise, beta is under investigation to inform DBS programming and adaptive DBS. This study assesses the applicability of these efforts by quantifying the biomarker’s prevalence and investigating its relationship to synchronized beta-bursting neurons.

**Methods:** STN LFP and spiking activity of n = 156 patients with Parkinson’s disease from six DBS centers recorded via microelectrodes was examined. Spectral peaks in the LFPs were classified and the absence of artifacts confirmed. The distribution of phase angles between beta LFP and spiking of individual beta-bursting neurons were explored.

**Findings:** Bilateral beta LFP expression was observed in 47·25% of patients, and in 65·59% of hemispheres. LFP-synchronized neuronal spiking clustered in two-thirds of neurons at a primary and in one-third at a secondary LFP phase (182·84° phase shifted). This applied on the group level and for individual patients across different DBS centers.

**Interpretation:** Beta-informed approaches mandate reliable biomarker expression, a requirement not met by slightly more than half of herein investigated patients. This is particularly troublesome for intraoperative guidance wherein the cause of biomarker absence, including DBS lead misplacement or a balanced ratio of phasic and antiphasic spiking neurons, cannot be determined, decreasing insight and increasing doubt.

Although beta LFPs remain a valuable biomarker, routine clinical practice should not rely on beta alone; instead, multiple biomarkers (i.e. beta-gamma coupling, finely tuned gamma and/or cortico-cortical beta-band coupling) should be evaluated simultaneously to increase robustness.

**Funding:** Maximilian Scherer received funding from the Alexander von Humboldt-Foundation. Thomas Koeglsperger has been supported by the Munich Clinician Scientist Program.

## Introduction

Increased subthalamic (STN) local field potential (LFP) oscillations in the beta frequency range (12-32 Hz) are considered a hallmark of Parkinson’s disease^1^, closely related to hypokinetic motor impairment. Suppression of STN-LFP beta via deep brain stimulation (DBS) accompanies symptomatic improvement^2^, prompting widespread adoption of beta as a biomarker to guide electrode implantation^3–8^, programming^9–11^, and adaptive DBS (aDBS)^12^. However, clinical performance of beta-informed strategies is inconsistent, and a substantial proportion of patients don’t express beta^3,9^. Reported per-hemisphere beta prevalence ranges from 64-69·5%^2,9,13^ in small cohorts to 84·8% (64·2% bilateral; n = 51) in the ADAPT-PD trial^14^ and 89·0% (69·4% bilateral; n = 113)^15^ in Medtronic’s BrainSense product surveillance registry (BrainSense PSR). Yet, the latter likely overstate true prevalence, as acknowledged by ADAPT-PD: First, beta was expanded to include alpha (8-12 Hz; both). Second, an undisclosed number of patients had been pre-screened for biomarker expression (ADAPT-PD). Third, peaks may have been defined as the *highest average power* (BrainSense PSR), intermingling beta peaks and background activity levels, which are higher inside compared to outside nuclei. Even under optimistic assumptions, beta-guided approaches may be inapplicable in at least one third of patients. Yet, without an unbiased prevalence estimate, the true generalizability of beta-based treatment approaches remains elusive.

The absence of beta has direct clinical implications. Electrophysiological mapping supplements anatomical targeting, which is limited by imaging resolution^16,17^ and brainshift^18^. Although beta LFPs offer a simpler alternative^19^ to 1998’s notoriously difficult to interpret^20^ gold-standard of microelectrode based guidance^21^, their utility depends on bilateral STN-LFP beta expression. While STN-LFP beta-based guidance is seemingly less complex, it introduces a strict reliance on bilateral biomarker expression. Nevertheless, 15·2-34·0% of implantations required revision or removal in North America from 2004 to 2013 with up to 48·5% potentially due to improper lead placement or insufficient therapeutic response^22^. Similar limitations affect guided DBS programming and aDBS: Beta informed approaches fail to identify clinically selected contacts in 8-50% of people^3,9,23^, show low co-variability (17·30%) with core motor symptoms, rigidity and bradykinesia^24^, and are not deployable in patients without beta expression^25^. Together with sporadic reports of the biomarker’s absence (n_pat_ = 3/8; n_hem_ = 7/19)^9,26^ and a strong publication bias (p < 0·006) favoring positive outcomes^27^, these observations suggest that beta prevalence may present a fundamental constraint to any beta informed system^12,28^.

The absence of STN-LFP beta in some patients remains unresolved. Beta LFPs have been shown to co-occur with beta-bursting neuronal spiking in the STN^1,21^ in humans, with spiking activity of beta bursting neurons^29^, particularly spiking within-bursts^30^ being synchronized to the LFP. However, pathological spiking and LFPs have been dissociated in animal models^31,32^ with spiking’s contribution outweighing oscillation’s impact on motor control^31^. While it is assumed that spiking may directly or indirectly contribute to LFP activity^33^, a causal explanation permitting beta-bursting neurons in the absence of detectable beta LFPs has not yet been established. One plausible mechanism is phase cancellation between subpopulations of beta-bursting neurons, which could locally preserve spiking synchrony while suppressing conglomerate phenomena such as beta LFPs.

Here, we quantify the prevalence of elevated STN-LFP beta power in 156 patients across six DBS centers and analyze single-neuron activity (606 units) to determine whether phase relationships among beta-bursting neurons account for the absence of detectable beta LFPs.

## Methods

### Patients

Data were pooled from publicly available sources^30,34,35^ and collaboration partners in Munich and Rostock, Germany, six DBS centers in total. Table 1 provides an overview of the employed data. Full details on data acquisition of publicly available data (cohorts 1-3) are provided in the respective manuscripts^30,34,35^ and the supplementary material (cohorts 1-5). In short, data were acquired either from externalized LFP leads between 1 and 6 days after surgery (cohort 1 & 2), or via microelectrodes in an intraoperative setting (cohorts 3, 4, and 5) and sampled between 2048 Hz - 24k Hz. Data were either unfiltered or applied filters were fully reversed (see supplementary figure 1). Full details on data origins, sampling, and filter reversal are provided in the supplementary material. Data distribution/local collection and evaluation have been approved by the respective ethics committees and written informed consent has been obtained from all study participants in accordance with the declaration of Helsinki.

### Anatomical reconstruction of microelectrode positions

Using computer tomography (CT) and magnetic resonance images (MRI) available, microelectrode trajectories explored in cohorts 4 and 5 were reconstructed, supporting their analysis. A reference trajectory was recovered via Lead-DBS^36^: The software was used to reconstruct the position of the implanted DBS lead in MNI space, localizing measurements along one microelectrode trajectory. The remaining microelectrode trajectories (up to 4; variable) were reconstructed via parallel projection matching the BenGun array’s shift (2mm), documentation on explored trajectories (i.e. central and lateral), and the intraoperatively employed ring and arc angles. Subsequently, measurement points along all microelectrode trajectories were determined by combining implantation depths, regularly spaced measurement points, and the structure of the implanted DBS leads. Reconstruction accuracy of DBS-lead positions, and microelectrode measurements was manually verified for each patient (see supplementary figure 2). 3D and 2D visualizations of the anatomical reconstructions of patients from cohort 4 and 5 are available in the supplementary material.

### Data acquisition and pre-processing

Data were collected from externalized DBS leads (cohorts 1 and 2; LFPs only) and microelectrodes (cohorts 3-5; LFPs and spiking). The data were zero-centered and detrended. A lowpass was applied (cohorts 1and 2: 299 Hz; cohorts 3/4/5: 4,999 Hz), and the data were downsampled (cohorts 1 and 2: 600 Hz; cohorts 3-5: 10,000 Hz). The data were manually screened for visually identifiable artifacts, (i.e. movement) and continuous, artifact-free recordings longer than 5 seconds (60 to 160 cycles of beta activity) were retained. Whenever multiple trajectory sets with different ring/arc angles were explored, only the final set with the DBS lead as reference point to recover anatomical position was evaluated.

### Beta peak detection

To identify beta peaks, the frequency spectra of all recordings obtained in a single hemisphere were concurrently visualized. Welch’s method (0·25-1 Hz frequency bin width) was employed to project signals into the frequency domain. Individual power spectra were visualized for 1-6, 6-14, 5-45, 10-36, and 15-36 Hz. Separate windows, normalized to their relative maxima, allowed for a highly resolved display, particularly at higher frequencies (see supplementary figure 3). A copy of each hemisphere’s LFP peak screening is provided in the supplementary materials.

### Quantifying LFP peak prevalence

Individual LFP peaks were identified and screened for artifacts. Analyses were conducted manually due to automated approaches inferior performance (30-76% accuracy) and major variance across subjects^37^. A detailed description on peak identification and artifact detection criteria, and a graphical documentation of each trajectory’s evaluation is provided in the supplementary material. In short, peaks were identified when LFP activity was locally elevated above noise levels in the alpha or beta bands. Given the availability of anatomical data for cohorts 4 & 5, peaks were additionally verified to be within the STN. Artifact screening included ensuring a variable amplitude across depths, a spectrally static peak, and the absence of low frequency harmonics. Leveraging cohorts’ 3 to 5 increased data density, non-reproducible peaks and spatially not limited peaks were labeled artifacts.

From these, we determined alpha and beta prevalence globally, per hemisphere, and per patient, the distribution of beta peak frequencies, and the ratio of encountered artifact types (i.e. non-repeatable or harmonics).

### Single-unit processing

The moving average of a recording was subtracted from the raw signal to delineate single unit activity (see supplementary figure 4). Subsequently, spikes were identified by applying a manually adjustable threshold. From these, bursts were delineated by identifying those spikes with an inter-spike-interval below a certain threshold, initialized as the average of within burst firing (∼10 ms^38^) and tonic activity rate (∼45 ms^39^), 20 ms. As spikes within bursts are more closely tied to the LFP ^30^, only those were retained. Via Welch’s method, the spectrum of the within-burst spiking activity was computed.

### Single-unit rating & analysis

To identify beta-bursting neurons, microelectrode recordings were screened for bursting neurons that expressed a beta spike in the neuron’s enveloped power spectrum. From these, we delineated recordings wherein bursting and low frequency peak were robust, expressed for an extended period and attributable to a single unit. These were termed *high fidelity* neurons. Akin to LFP evaluations, a full description of assessment criteria and assessments of all investigated single neurons are available in the supplementary material. Any analyses were performed on both, all and high fidelity neurons exclusively.

Phase amplitude coupling (PAC; direct modulation index^40^) and coherence (directional absolute coherence^41^) between the within burst spiking activity and the LFP were calculated, the latter filtered to the bursting peak frequency ±3 Hz. Magnitude and phase were extracted from the PAC estimate. Magnitude and directionality from the coherence estimate. Neurons with a PAC of < 0.1 (scale: 0 to 1) were discarded.

### Data and code availability

Data from cohorts 1-3 is publicly available^30,34,35^. Data from cohorts 4 & 5 may be shared upon request. Python code written to support this evaluation has been made available at: https://codeberg.org/MaximilianScherer/2026_MS_beta_prevalence.git^42^. Additional supplementary material (LFP/spike evaluations and microelectrode reconstructions) is available at zenodo^43^.

## Results

### Bilateral prevalence of STN-LFP beta power is 47·25% per patient

To analyze the prevalence of elevated STN-LFP beta power, data acquired from extra- (cohorts 1 and 2) and intraoperative recordings (cohorts 3-5) were analyzed. Without artifact rejection, per-hemisphere prevalence was 74·54% (average; see Figure 1A: partially transparent *low insight* bars). Post artifact rejection, per-hemisphere STN-LFP beta prevalence was 65·59% (average; see Figure 1A: colored *high insight* bars). These results were mostly consistent across all cohorts, with cohort 1 having the highest and cohort 2 the lowest prevalence. Bilateral prevalence (left and right hemisphere) of STN-LFP beta was 47·25% per patient (average; see Figure 1B) when rejecting artifacts 54·94% without artifact rejection. Unilateral prevalence (left or right hemisphere, or both) of STN-LFP beta without artifact rejection was 90·10% (average; see Figure 1C), 83·51% after artifact rejection. Comparing STN-LFP beta power prevalence between the left (Figure 1D) and right hemisphere (Figure 1E) did not reveal any systematic differences. Of note, as data from cohort 3 could not be attributed toward a specific hemisphere, data is visualized in left-hemisphere plots (see Figure 1A/D). Any beta peak candidates identified as artifacts were classified as such for the following reasons: Beta peaks were post-hoc localized as not within the STN (63·93%; cohorts 4 & 5 only; see Figure 1F). They were not reproducible (16·39%; cohorts 3-5 only). They were another type of artifact, including persistent effects and harmonics (19·67%).

**Figure 1:**
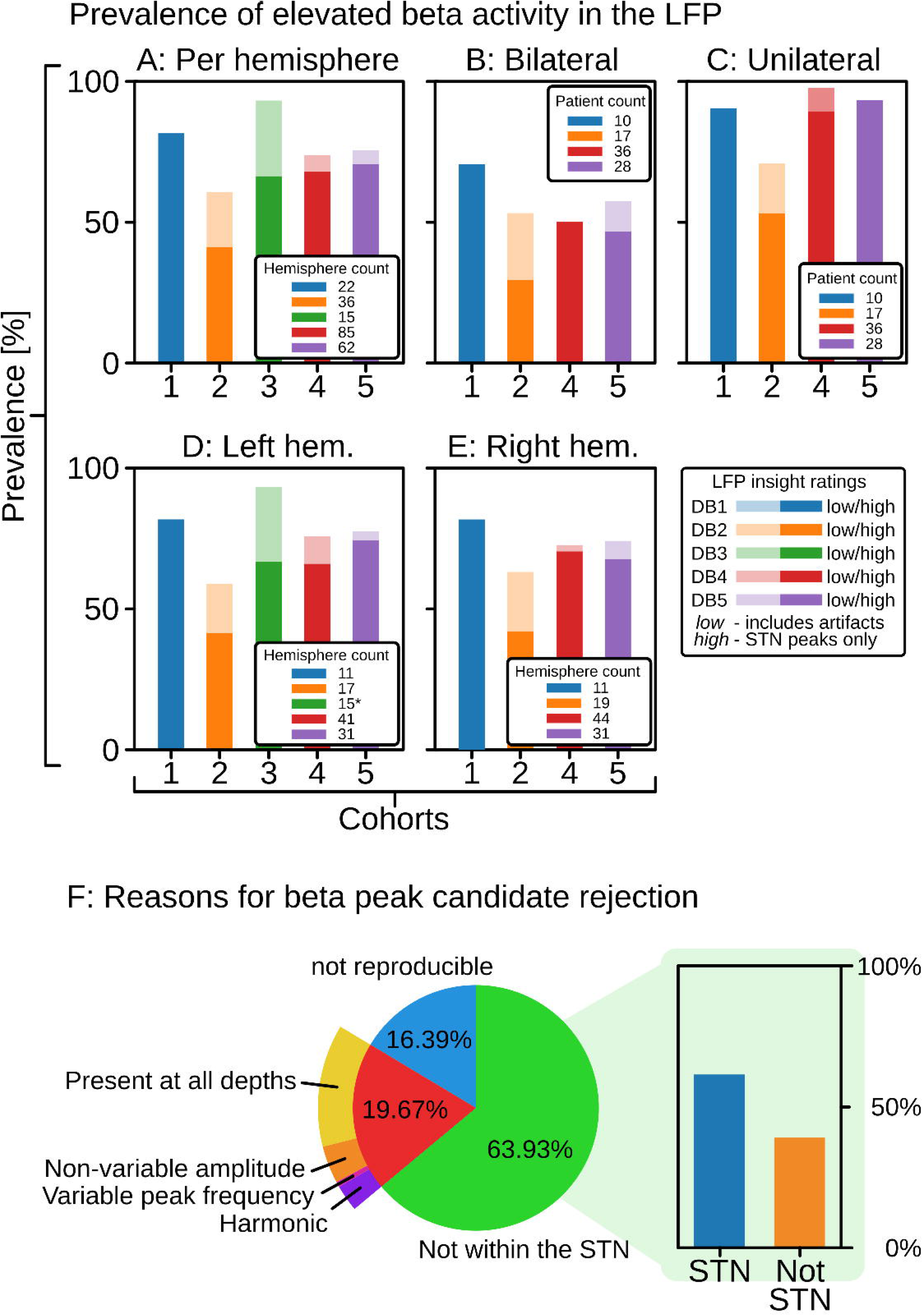
Prevalence of elevated beta band local field potentials (LFPs) in the subthalamic nucleus (STN) of people with Parkinson’s disease, per hemisphere (A; 65·59%), bilaterally (B; 47·25%), unilaterally (C; 83·51%), in the left hemisphere (D; 66·08%), and the right hemisphere (E; 65·71%), separated into counts without (low insight; partially transparent) and with (high insight; opaque) artifact rejection. Beta peak candidates were rejected (F) for being outside the STN (63·93%), being not reproducible (16·39%), or another type of artifact (16·67%)

Finally, alpha activity was less prevalent (as expected), presenting a far more sporadic expression (See supplementary figure 5).

### Beta bursting neurons are phase locked to STN-LFP activity at two opposing angles

Having previously observed a strong coupling between within burst spiking activity of single neurons and LFPs^30^, this relationship was further explored by quantifying the phase angle homogeneity between the two electrophysiological phenomena. Including all neurons, activity synchronized to the LFP at 234·50° (Figure 2A) with a secondary population at −20·18°, yielding a phase difference of 254·68°. When focusing on high fidelity, high PAC neurons, peaks shifted to 235·19° and 52·35°, with a phase difference of 182·84°. This observation was replicated across individual patients across all applicable cohorts (3, 4, and 5; Figure 2C & D). Calculating an artificial composite signal of each patient’s neurons demonstrated that phase shifted neurons resulted in low magnitude conglomerate/multiunit activity (Figure 2C) whereas phase synchronized ones resulted in high magnitude conglomerate activity (Figure 2D).

**Figure 2:**
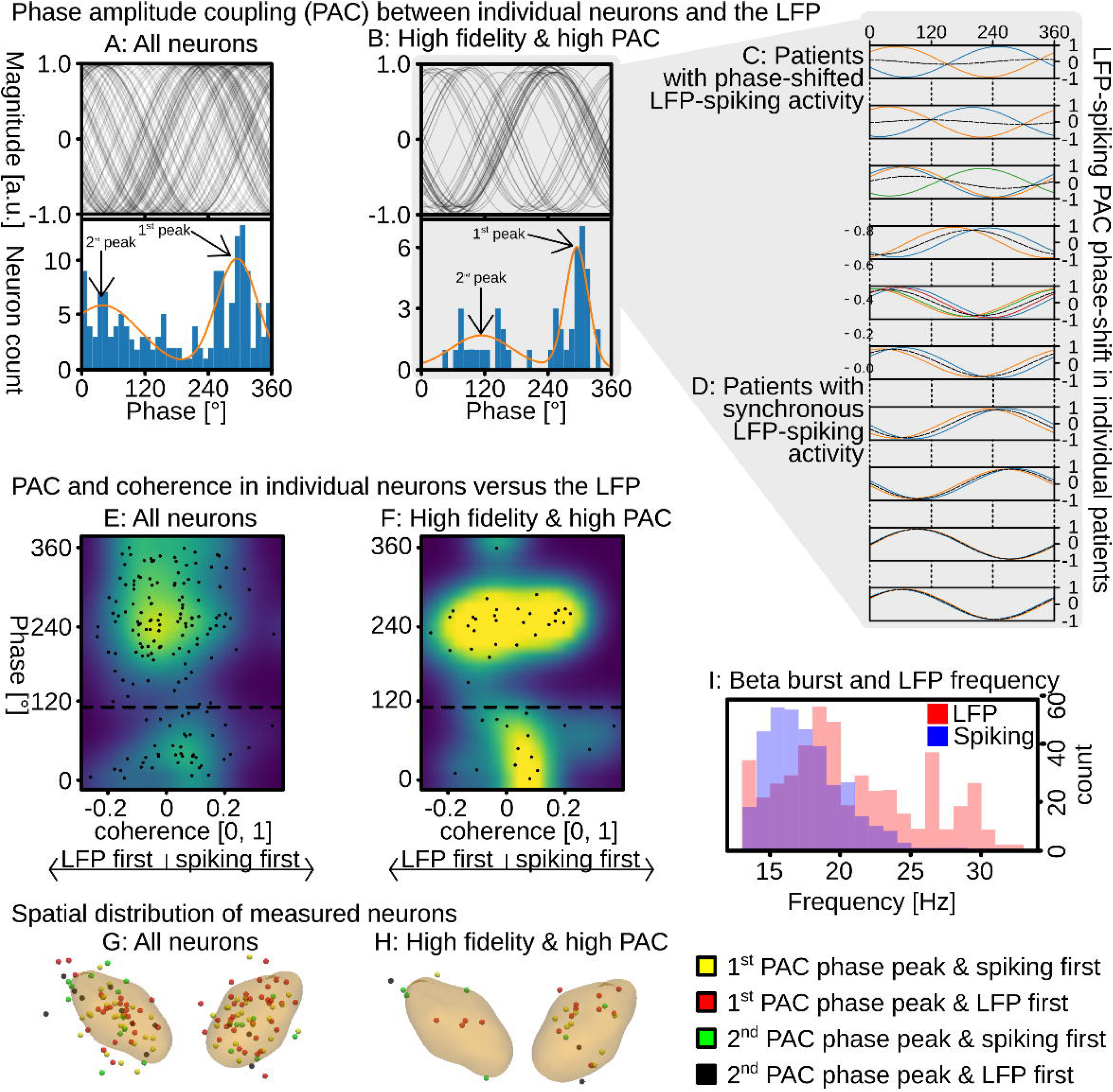
Phase amplitude coupling (PAC) between within burst neuronal spiking aligned with the LFP at a phase of 234·50° for all neurons (A) and 235·19° (B) for high fidelity and high PAC neurons. A 2^nd^ phase cluster (peak) was identified between −20·18° and 52·35°, yielding a phase difference of 182·84° to 254·68°. The observation of different spiking to LFP phase alignments was replicated in some (C), but not all patients (D). The estimated conglomerate activity is indicated via a dashed black line. Further analyses revealed a difference in coherence between these two populations of neurons. Namely, neurons clustering at the 1^st^ peak did not exhibit a significant preference whether the spiking or LFP occurred first (E & F – upper), yet spiking of the 2^nd^ peak population mostly preceded the LFP (E & F – lower). A spatial difference between these two populations of neurons was not observed (G & H). Beta peaks for enveloped spiking activity and LFP were similar (I).

Further analysis revealed that these populations did not only differ in their phase angle to the LFP, but also their coherence to the LFP. Namely, while there was no distinct causal sequence (p < 0.06; 56 vs. 64 neurons) between the LFP and the enveloped bursting signal of 1^st^ peak neurons (234·68°; Figure 2E, upper panel), neuronal spiking did precede (p < 0.02; 15 vs. 29 neurons) the LFP for the 2^nd^ peak (−20·18°; Figure 2E, lower panel). Similarly, there was no distinct causal sequence (p < 0.12; 13 vs. 17 neurons) when focusing only on the enveloped bursting signal of high fidelity, high PAC 1^st^ peak neurons (235·19°; Figure 2F, upper). Yet, neuronal spiking again preceded (p < .0418; 4 vs. 11 neurons) the LFP for the 2^nd^ peak (52·35°; 2F, upper). Analyzing the spatial distribution of these neurons (Figures 2G & 2H) did not reveal any succinct patterns.

Finally, comparing frequency distributions of the LFP and spiking activity revealed similar peaks (Figure 2I). Calculating an artificial composite signal of all neurons PAC curves confirmed that the amplitude of the neuron’s summed activity was lower if neurons were antiphasic and higher, if the neurons were in-phase (see Figure 2E and F: black striped lines).

## Discussion

Electrophysiological analyses are increasingly employed in daily medical routine, however, the prevalence of underlying biomarkers often remains insufficiently investigated. Herein, we explored the prevalence of subthalamic (STN) local field potentials (LFPs) in the beta frequency band (13-32 Hz), considered a hallmark in people with Parkinson’s disease^1^. This biomarker is used to guide DBS lead implantation, programming, and adaptive stimulation, all of which require reliable bilateral biomarker expression to be applicable or perform optimally. However, herein we found that this biomarker is only expressed in slightly less than every other patient bilaterally (47·25 %).

Exploring the relationship between neuronal spiking activity and LFPs provided a potential explanation. Namely, we observed two distinct populations of neurons, being coupled to the LFP at 52·35° and 235·19°. While the exact relationship between spiking and LFP remains elusive, the herein observed partially phasic-antiphasic relationship of neurons to the LFP may be disadvantageous for LFP production. Furthermore, these two populations also expressed varying causal relationships to the LFP. Namely, the secondary population was significantly tilted toward acting prior to the LFP whereas the main population did not express any observable preference, supporting the identification as two separate populations.

### Prevalence of STN-LFP beta power in the STN

The ratio of bilateral STN-LFP beta expression observed in each cohort investigated herein is lower than those reported in both the ADAPT-PD trial (64·0 %)^14^ and the BrainSense PSR (69·4%)^15^, at 47·25%. Likewise, while per-hemisphere prevalences are equally lower (65·59% vs 84·8% and 89·0%, respectively), they matched previous small scale reports^2,9,13^ (64-69·5% vs. 65·26%). Varying definitions of beta activity may have contributed to this difference with the ADAPT-PD and PSR trials including alpha (8-12 Hz) in the beta range. However, given the largely inconsistent pattern of alpha activity observed herein, this aspect is unlikely to be the sole cause. An alternative explanation may lie in trial specific differences. Namely, ADAPT-PD prescreened an undisclosed number of patients for beta and PSR potentially treated the contact with the highest average power as a beta peak. Beyond these systematic differences, extra-subthalamic beta peaks may have also been inadvertently misclassified as STN-LFP beta. The observation of such beta peaks in the substantia nigra pars reticulata (SNr) was common herein and in previous reports^44,45^. Hence, beta peaks, particularly at the end of DBS electrodes or in deeper implanted DBS leads may have erroneously been misclassified in cases where not all electrode positions had been post-hoc reconstructed to verify proper positioning. Finally, approximately one in six beta peaks observed herein were not reproducible as these appeared in a single measurement and never thereafter. Hence, it is conceivable that noise-like artifacts may have inflated beta prevalence further.

### STN-LFP beta power guiding DBS lead implantations

The region of strongest beta elevation has repeatedly been observed to co-localize with the optimal DBS lead implantation site within the STN^3,46^ and correlate significantly with symptoms^20^. As, post-hoc analyses have demonstrated, the optimal implantation trajectory is often marked by strong beta expression^3^, in up to 80% of cases^47^, the biomarker has been suggested as a reliable indicator of proper lead placement^48^. Conversely, beta activity was only observed in 65·59% (vs. 64-69·5% small scale reports^9,13,49^) of hemispheres with bilateral prevalence at only 47·25%, slightly less than every other patient. While the above discussed causes may technically provide an answer to this discrepancy (i.e. SNr beta and/or increased broadband activity within nuclei), intraoperative information is usually too limited to determine the cause of biomarker (non-)expression. Namely, an increase of beta LFPs may also be observed when entering any nucleus (increase in broadband power), entering the STN or SNr specifically, or due to noise forming a non-reproducible beta peak. Likewise, absence of beta LFPs may be observed when misplacing DBS leads or in patients without biomarker non-expression (34·41% of hemispheres). Hence, it may be difficult to select the optimal trajectory from beta LFPs alone. While beta LFPs may provide important additional insights, it is advisable to corroborate these with microelectrode recordings and/or imaging information.

### STN-LFP beta power in DBS programming and adaptive DBS

STN-LFP beta is considered a reliable biomarker^48^ to assess DBS related treatment benefits^2^. As such, it has been selected to guide several manual and (half-)automated DBS programming (i.e. Medtronic BrainSense^15^), and adaptive DBS (aDBS) approaches. Yet, while significantly correlating with symptom expression^20^, LFP variability only poorly captures symptom severity, indicated by qualitatively weak correlation (r² ≈ 17%)^24^. Correspondingly, while initial assessments of beta informed aDBS had been very positive, albeit with a significant publication bias toward positive outcomes^27^, recent trials’ outcomes are more modest^50^, with improved ON-time over continuous DBS in 36 cases (52·94%) and no impact or worsened in 32 cases (47·06%)^51^. The often exclusive use of beta^12^ limits applicability to patients with at least unilateral biomarker expression, as demonstrated by Oehrn et al.^25^. Yet, it is assumed that bilateral sensing is required for optimal aDBS performance^52^. However, this condition is not met by 52·75% of patients. A potential way forward may lie in diversification, levering a set of biomarkers including beta-gamma coupling^25,53^, finely tuned gamma oscillations^34^ and/or cortico-cortical beta-band coupling^54^. While individual biomarkers may not be reliably expressed in every patient, concurrent non-expression of all biomarkers in a larger set is assumed to be less likely.

### Beta-bursting neurons may mask STN-LFP beta power

While the exact relationship between single neuron and conglomerate multi-unit activity, such as LFPs, has not yet been determined, it is assumed that LFPs may constitute at least in part from synaptic currents and synchronized spiking activity^33^. Spiking activity has been shown to synchronize at a specific phase of the LFP^29^, especially when occurring within bursts^30^. Herein, this concept was expanded by characterizing the phase-angle between spiking and LFP activity. While most neurons’ spiking synchronized at 235·19°, with one-third of neurons’ spiking synchronized at 52·35°, a 182·84° phase shift. This result was repeated across individual patients from all three applicable cohorts investigated herein. Conceptually, activity of this antiphasic minority may dampen overall LFP activity levels which are measured by a macroelectrode or sensed from a DBS lead. Namely, as contributions from individual neurons likely average in multi-unit LFP activity ^33^, individual neuron’s activity must be aligned to produce a high magnitude LFP. However, if phasic and antiphasic, the sum of individual neuron’s activity would negate on a macroscopic scale, with a continuous effect on the LFP magnitude between aligned and phasic/antiphasic scenarios. While the ratio of in-phase and antiphasic neurons within an individual human subject cannot be determined at this point, the herein established concurrent presence of neurons from both populations within the same patients provide a causal explanation for reduced or absent beta activity in the STN-LFP – given the assumption that spiking significantly contributes to the LFP^33^.

## Limitations

Low-frequency attenuation introduced by low order hardware filters in cohorts 4 and 5 was reversed through inverse filtering to recover the raw signals. Albeit the application of a filter is a simple multiplication in the frequency domain, an easily reservable step, this may have introduced additional noise, as does most data processing. However, the original signal was numerically stable, and correct filter reversal was manually confirmed (see supplementary figure 1) supported by the close similarity of results across cohorts. For cohorts 4 and 5, only the final implantation trajectory was analyzed in patients with multiple trajectories. While DBS lead positioning and microelectrode trajectory projections were verified via anatomical reconstruction and visual confirmation, anatomical inaccuracies may have impacted beta estimates. To compensate, beta peaks within 1 mm of the STN were still considered inside the STN. While anesthesia can influence beta activity ^55^, no recordings were obtained during active anesthesia. Cohort 3 was acquired under local anesthesia, whereas cohorts 4 and 5 were obtained predominantly under general anesthesia that was discontinued at least 10 minutes before recording; nevertheless, bilateral and unilateral beta prevalence was comparable across centers. Postoperative microlesion effects^56^ may have increased noise^57^ in cohorts 1 and 2, but their combined results remained consistent with intraoperative cohorts (3 to 5).

## Conclusion

Albeit bilateral STN-LFP beta expression was observed in only 47·25% of patients, the biomarker offers valuable information that can be leveraged for patient benefit. However, given its limited prevalence, additional biomarkers such as beta-gamma coupling, finely tuned gamma, and/or cortico-cortical beta-band coupling may be assessed in parallel to increase robustness. This approach allows for reliable estimates – such as confirming proper DBS lead placement when a single biomarker is not expressed.

## Conflicts of interest concerning the research related to the manuscript

Thomas Koeglsperger has received research funding from Medtronic and Abbott and speaker honoraria from Abbott and AbbVie. Thomas Koeglsperger and René Reese serve as the president and vice-president of the German DBS Society (AG-THS).

