## Supplementary material for "Inconsistent Subthalamic Beta Expression in the Local Field Potential amid In- and Anti-Phasic Neuronal Bursts"

### Supplementary Materials

#### Methods

##### *Data set/cohort descriptions*

<u>Dataset/cohort 1</u>

This dataset comprises multi-center recordings from externalized patients (n = 16), acquired 3-6 days after DBS lead implantation into the STN[27]. The data were sampled at 2048 Hz via a TMSi Porti (TMS International, Netherlands) or at 4096 Hz via a TMSi Saga32 (TMS International, Netherlands). Activity was recorded from the implanted DBS lead's ring contacts. These were treated as measurements at different depths herein. This analysis only included baseline data.

<u>Dataset/cohort 2</u>

This dataset comprises recordings from externalized patients (n = 19), acquired one day after DBS lead implantation in Düsseldorf, Germany[28]. The data were sampled at 2 kHz via a custom connected MEG system (VectorView, MEGIN), and bandpass filtered between 0·1 and 660 Hz. Activity was recorded from the implanted DBS lead's ring contacts. These were treated as measurements at different depths herein. Only resting state data during med OFF was used in this investigation.

<u>Dataset/cohort 3</u>

This dataset comprises intraoperatively acquired recordings (n = 15) from Toronto, Canada, collected while mapping the STN's extend using two microelectrodes[23] in a custom setup. The data were sampled at ≥12·5kHz via two Guideline System GS3000 amplifiers (Axon Instruments, USA). Measurements were taken along the implantation trajectory, whenever a neuron was clearly visible.

<u>Dataset/cohort 4</u>

This dataset 4 comprises intraoperatively acquired recordings (n = 67) from Munich, Germany, collected while mapping the STN's extend using between 1-3 microelectrodes in a Ben-Gun array. The data were sampled at 20kHz via an Isis IOM system from Inomed, Germany. Measurements were taken every 0·5 mm, starting 5 mm before the STN's center for at least twelve consecutive steps. Activity outside 30-5,000 Hz was mildly attenuated by a low-order hardware filter. As filtered frequencies remained above floating point precision and exact filter knowledge was obtained, the filter's application was fully reversible, restoring raw, unfiltered, data. Successful signal recovery was manually confirmed, see Supplementary Figure 1: A1 and A2. For more details, see Supplementary Materials.

<u>Dataset/cohort 5</u>

This dataset 5 comprises intraoperatively acquired recordings (n = 39) from Rostock, Germany, collected while mapping the STN's extend using 3-5 microelectrodes in Ben-Gun array. The data were sampled at 24 kHz via a Leadpoint system from Natus, Germany. Measurements were taken every 1 mm starting 5 mm before the STN's center for ten consecutive steps. Activity outside 500-5,000 Hz was mildly attenuated by a low-order hardware filter. As filtered frequencies remained above floating point precision and exact filter knowledge was reverse-engineered, the filter's application was fully reversible, restoring raw, unfiltered, data. Successful signal recovery was manually confirmed, see Supplementary Figure 1: B1 and B2.

***Signal recovery/Hardware filter removal***
<u>*Dataset 4*</u>
Acquired signals were processed by an RC (resistor-capacitor) filter, producing frequency-dependent signal attenuation and phase distortions below 30 Hz. Filter-parameters were as follows, r = 5.6kOhm and c = 1µF. As the implemented hardware filter is a 1st order filter, (having only one degree of freedom, the rolloff), it can be digitally implemented via a Butterworth filter. Hence, the application of the RC filter can be post-hoc reversed by applying an inverted digital clone of the RC filter. Said inversion may be achieved by swapping the filter's zeros and poles. Thereby, frequency-dependent signal attenuation is undone, as is the filter induced phase distortion. Afterward, proper signal recovery was confirmed via manual inspection (see Supplementary Figure 1: A1 and A2).

<u>*Dataset 5*</u>
Acquired signals were processed by a hardware filter with a cutoff at 500 Hz. Reverse-engineering identified the filter as a 1st order filter. As 1st order filters are defined by a 3 dB attenuation at the cutoff frequency as the sole parameter, a digital clone was created using a 1st order Butterworth filter, sharing this property. Applying the clone with reversed zeros and poles restored the original signal. Proper signal recovery was confirmed via manual inspection (see see Supplementary Figure 1: B1 and B2).

***Filter reversals applied to dataset/cohort 4 & 5***
Data from datasets/cohorts 4 & 5 were recorded with low order hardware filters enabled. Due to the employed filter's low order (with -3 db at the cutoff; signal strength reduced to ~70%), information was not pushed beyond numerical accuracy of the respective employed float types. Hence, by applying the inverse of the originally employed filters, the original amplitudes and phases were recovered. This is demonstrated in Figure 1 for dataset 4 (upper) and dataset 5 (lower). After filter reversal (orange), the 1/f gradient (pink noise) is fully restored.

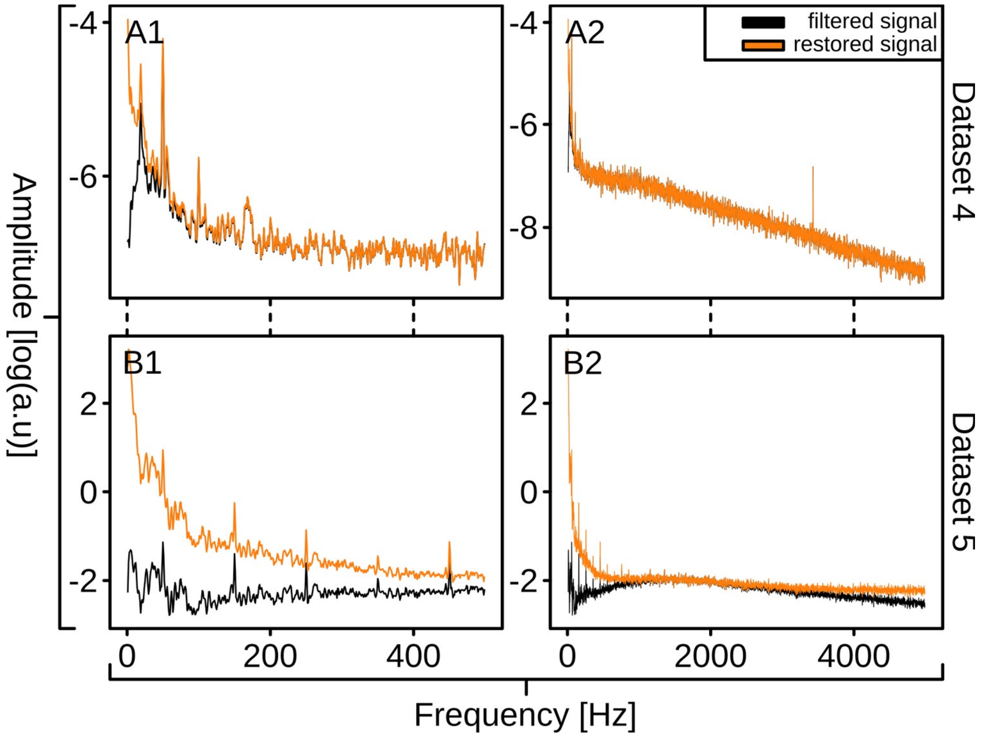

*Supplementary Figure 1: Frequency domain signals prior (black) and after (orange) filter reversal. Pink noise (1/f) is restored, as is information in the corresponding frequencies. A1 and A2 show a sample for database 4 and B1 and B2 for database 5.*

***Visualizations of reconstructed microelectrode trajectories***

DBS-lead implantation positions were reconstructed via lead-DBS. Leveraging information on the selected implantation trajectory, depth, and implanted electrode type, in combination with the intraoperatively employed ring and arc angles, and explored microelectrode trajectories (i.e. lateral or anterior), microelectrode trajectories were projected into 3D space. These projections were manually verified for all patients of cohorts 4 and 5. A sample is provided below (see Supplementary Figure 2). Reconstructed trajectories for all patients may be found in the downloaded zip file:

- "*microelectrode_reconstructions/database4*"
- "*microelectrode_reconstructions/database5*"

These contain 2D illustrations as PNG files and drag and drop enabled 3D visualizations as HTML files. The latter can be opened with any (web) browser.

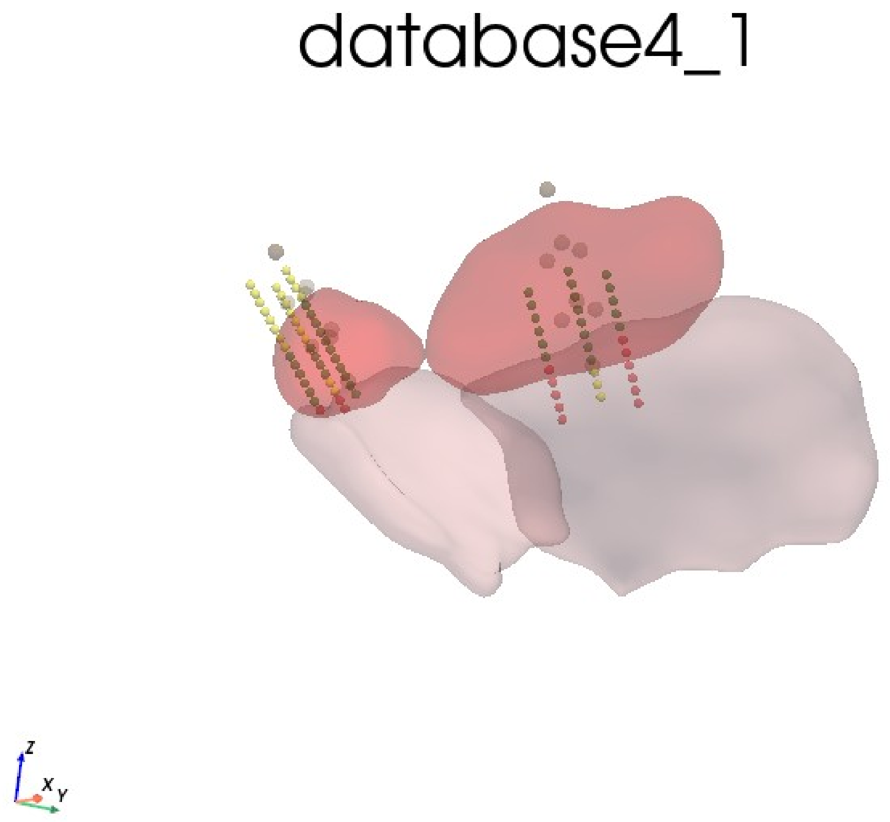

*Supplementary Figure 2: Sample 2D visualization of explored microelectrode trajectories from the 1st patient in dataset/cohort 4*

***LFP detection tool***
Sample of the custom written LFP peak detection tool employed in this investigation.

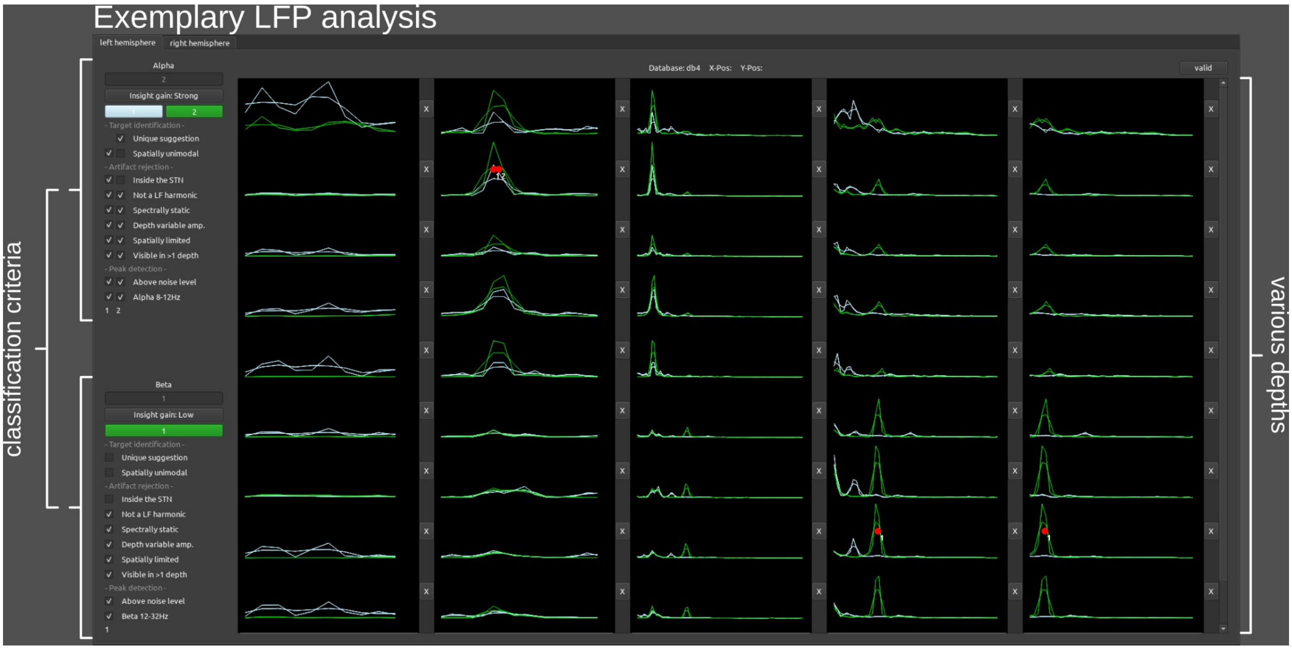

*Supplementary Figure 3: Custom UI developed to identify beta LFP peaks.*

Results for all investigated patients are available in the provided ZIP file in the following directories:

- "*lfp_peak_detection/database1/*"
- "*lfp_peak_detection/database2/*"
- "*lfp_peak_detection/database3/*"
- "*lfp_peak_detection/database4/*"
- "*lfp_peak_detection/database5/*"

Criteria for LFP peak selection are described in the following paragraph.

***LFP detection criteria***
The requirements for peak detection and artifact rejection are described below.
<u>*Peak detection*</u>
*Requirements:* An LFP peak had to be in the range of 8-12 Hz or 12-32 Hz, above the noise level observed in the signal, and peaks additionally had to be within the STN (cohorts 4 and 5 only).
*Example:* A peak at 15 or 20 Hz would meet this requirement, if its elevation was beyond the noise level in the investigated signal.

<u>*Artifact rejection*</u>
*Requirements:* Be not a harmonic, be spectrally mostly static, and express a depth variable amplitude. Via these criteria, we separated disease-specific peaks in spectral activity from electrical artifacts (i.e. line noise – constant amplitude/spatially unlimited), other artifacts (i.e. harmonics). Furthermore, leveraging the increased data density of cohorts 3 to 5, peaks therein also had to be spatially limited, and visible across multiple recordings. *Example:* In case a peak's 15 Hz activity was mirrored by activity at 10 Hz and 5 Hz, it was classified as a harmonic distortion. In case it would have been expressed across too many investigated depths (> 8 mm), it would fail to meet the spatially limited requirement. Finally, in case the peak was only observed in a single measurement and not within 2 mm thereafter, it was considered non-reproducible and hence not disease-specific.

***Bursting activity classification tool***

Sample of the custom written spiking and burst detection tool employed in this investigation.

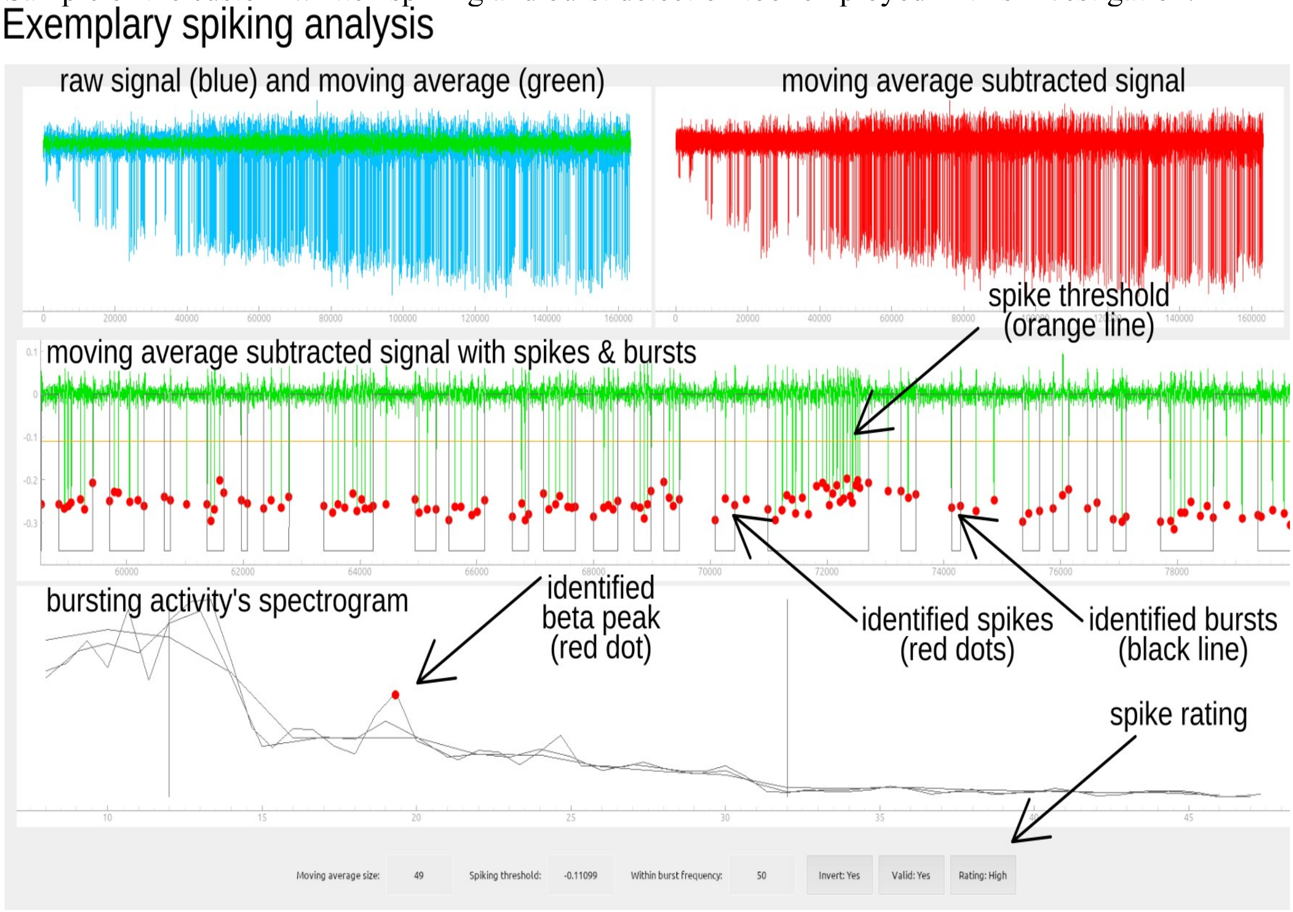

*Supplementary Figure 4: Custom UI developed to identify beta bursting neurons.*

Results for all positively screened recordings are available in the provided ZIP file in the following directories:

- "*spike_burst_detection/database1/*"
- "*spike_burst_detection/database2/*"
- "*spike_burst_detection/database3/*"
- "*spike_burst_detection/database4/*"
- "*spike_burst_detection/database5/*"

Criteria for spiking and burst detection are described in the following paragraph.

***Microelectrode recording screenings***

Microelectrode recordings were screened for visibly observable single unit activity. Once detected, a moving average filter was employed to identify and subtract the low-frequency component from the signal. Subsequently, a threshold was employed to isolate individual spikes. Afterward, a threshold was set to capture clustering of individual bursts. Finally, the power spectrum of the selected enveloped spiking activity was computed and investigated for the presence of a beta frequency peak. Peaks in the enveloped spiking activity were only marked as such if these were robust toward minor adjustments in analysis parameters (moving average size, spiking threshold, and within burst frequency).

***Single neuron ratings***

Neurons were labeled high fidelity, if their activity met the following criteria: Activity was attributable to a single unit, expressed a robust bursting pattern over an extended period of time, as well as a robust peak in the enveloped activity spectrogram. For example, in case a recording

comprised the activity of multiple units, that neuron's activity was not labeled high fidelity, if a separation between the primary and other units was possible but difficult. However, if the separation was not difficult, it was rated high fidelity.
Neurons were labeled high PAC if the PAC between spiking activity and enveloped spiking signal did exceed a threshold of 0.4 on a scale from 0 (no PAC) to 1 (full PAC).

***Individual LFP and spike assessments, and microelectrode reconstructions***
These are available on zenodo (doi.org/10.5281/zenodo.17951551).

#### Results

##### *Alpha power prevalence*

Peaks in alpha power (Figure 5), the control condition, were far less prevalent, as anticipated. Furthermore, variability and artifact presence were higher as well.

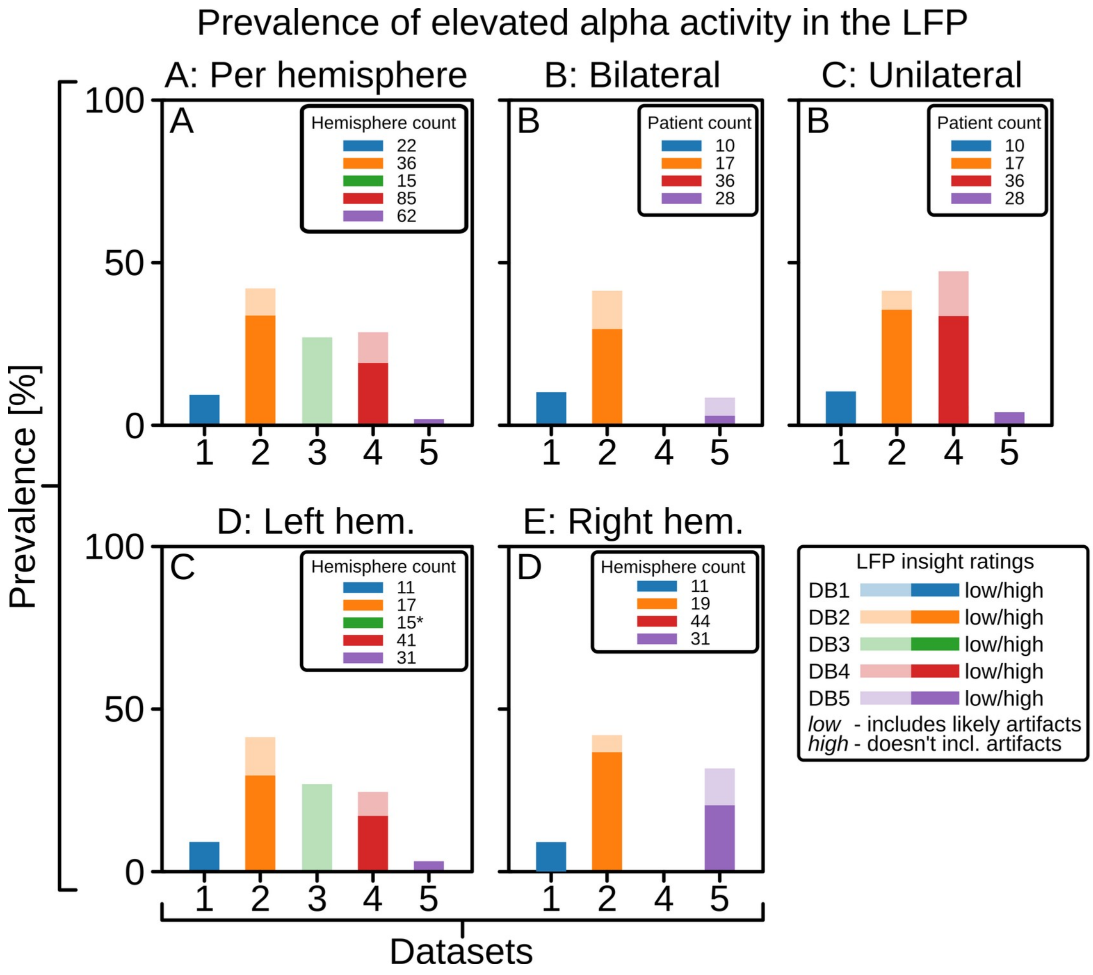

*Supplementary Figure 5: Alpha power peak prevalence.*
